# Rodent-driven NO_3_^−^-N enrichment reshapes amoeba–bacteria co-occurrence and bacterial functional potential in burrow soils

**DOI:** 10.64898/2026.04.30.721900

**Authors:** Chunfang Zhang, Florent Sebbane, Chutian Zhang, Jason D. Whittington, Yong Zhao, Chaolemen, Ruifu Yang, Nils Christian Stenseth, Lei Xu

**Affiliations:** School of Life Sciences, Zhengzhou University, Zhengzhou, Henan 450001, China; Centre for Ecological and Evolutionary Synthesis, Department of Biosciences, Faculty of Mathematics and Natural Sciences, University of Oslo, Oslo N-0315, Norway; University of Lille, Inserm, CNRS, CHU Lille, Institut Pasteur de Lille, U1019—UMR9017—CIIL—Center for Infection and Immunity of Lille, F-59000 Lille, France; Vanke School of Public Health, Tsinghua University, Beijing 100084, China; Institute for Healthy China, Tsinghua University, Beijing 100084, China; College of Natural Resources and Environment, Northwest A&F University, Yangling, Shaanxi 712100, China; Centre for Pandemics and One-Health Research, Sustainable Health Unit, Institute of Health and Society, Faculty of Medicine, University of Oslo, Oslo N-0316, Norway; State Key Laboratory of Pathogen and Biosecurity, Academy of Military Medical Sciences, Beijing 100071, China; Chen Barhu Banner Center for Disease Prevention and Control, Hulunbuir, Inner Mongolia 021599, China

**Keywords:** Rodent burrow, Soil factor, Amoeba–bacteria interactions, Pathogenicity-related functions, Nitrogen cycling

## Abstract

Interactions between amoebae and bacteria are increasingly viewed as key drivers of zoonotic pathogen emergence in rodent-dwelling burrows, yet the environmental factors shaping these interactions remain poorly understood. Here, we analyzed soil characteristics and used absolute quantitative high-throughput sequencing to assess microbial communities in active burrow, inactive burrow, and off-burrow soils across four rodent species (marmot, squirrel, gerbil, and vole) in the Hulunbuir grassland of Inner Mongolia, China. This study demonstrates that rodent activity creates chemically distinct soil microhabitats, with nitrate (NO ^−^-N) enrichment in active burrow soils consistently observed across rodent species. Elevated soil NO_3_^−^-N was associated with reduced microbial phylogenetic diversity and reorganization of amoeba-co-occurring bacterial assemblages. Both absolute abundance-based correlations and functional prediction of co-occurring bacteria indicated that amoebae were primarily associated with nitrogen-cycling bacteria in off-burrow soils. In burrow soils, amoebae increasingly interacted with bacterial taxa associated with pathogenicity while retaining ties to nitrogen-cycling taxa. Structural equation modeling and mediation analysis revealed that NO_3_^−^-N enrichment indirectly linked to increased infectious disease-related functional potential by amoeba-associated bacterial restructuring and coordinated shifts in nitrogen cycling, independent of changes in bacterial abundance. Together, our findings highlight the importance of rodent-driven soil heterogeneity in shaping amoeba–bacteria interactions and suggest that rodent-mediated NO ^−^-N enrichment may promote the emergence and persistence of potentially pathogenic bacteria, with broader implications for soil ecosystem functioning and disease-related processes in terrestrial ecosystems.

## 1. Introduction

Rodents act as ecosystem engineers, generating fine-scale heterogeneity in soil environments through burrowing and nutrient redistribution (Davidson and Lightfoot, 2008; Mallen-Cooper et al., 2019). These activities create localized soil microhabitats within burrows that differ substantially from off-burrow soils (Cai et al., 2026; Clark et al., 2016). However, empirical evidence suggests that many soil properties respond to rodent activity in a context- or species-dependent manner (Huang et al., 2023; Mallen-Cooper et al., 2019), raising the question of whether any soil factors exhibit consistent responses across rodent species, and how such consistency may structure soil microbial communities.

Rodent-induced soil disturbance has been increasingly recognized as an important driver of microbial diversity, yet the underlying mechanisms remain poorly understood (Dong et al., 2024; Xiong et al., 2025). Nitrogen enrichment in burrow soils, resulting from rodent excreta and enhanced mineralization, can modify microbial community assembly by changing resource availability, imposing environmental filtering, or mediating biotic interactions among microorganisms (Gao et al., 2025; Goberna et al., 2014; Horner-Devine and Bohannan, 2006). These processes can influence both taxonomic composition and phylogenetic structure, favoring taxa adapted to nitrogen-rich conditions while potentially reshaping microbial coexistence and associated functional capacities (Gao et al., 2025; Goberna et al., 2014). However, how such nitrogen-driven changes in soil conditions influence microbial interactions and shape microbial diversity and functional potential in rodent burrow soils remains largely unknown.

Free-living amoebae are ecologically important yet understudied microbial predators within soil food webs and have been proposed as potential environmental reservoirs for bacterial pathogens (Molmeret et al., 2005; Shi et al., 2021). By preying on bacteria, amoebae impose strong selective pressures that favor traits related to intracellular survival and stress tolerance, some of which overlap with mechanisms associated with pathogenicity (Jules and Buchrieser, 2007; Markman et al., 2018). Nitrogen availability and nitrogen cycling processes can modulate amoeba–bacteria interactions in soils (Cecil and Yoder-Himes, 2024; Elliott et al., 1979), suggesting that rodent-driven modifications of burrow soils may indirectly shape the composition and functional potential of amoeba-associated bacterial assemblages. However, how rodent-mediated soil heterogeneity influences amoeba–bacteria interactions and functional implications in soil ecosystems remains largely unexplored.

Here, we investigated whether rodent activity creates chemically distinct soil microhabitats along a burrow-to-off-burrow gradient that reshape amoeba-co-occurring bacterial assemblages through nitrogen-driven changes in soil conditions and associated microbial interactions, ultimately influencing microbial functional potential.

## 2. Materials and methods

### 2.1. Soil sampling, plant survey, and determination of soil characteristics

Soil samples were collected in the Hulunbuir grassland of Inner Mongolia, China (Fig. 1). The sampling map was created using ArcGIS (version 10.8). We sampled active burrow, inactive burrow, and off-burrow soils across habitats of *Marmota sibirica* (Marmota), *Spermophilus dauricus* (squirrel), *Meriones unguiculatus* (gerbil) and *Lasiopodomys brandtii* (vole), with three replicates per site (Fig. S1). Burrow wall soils were sampled at 0.5 m depth along the burrow path using soil corers with minimal disturbance. Off-burrow soil samples were collected from the same landscape at least 500 m away from both active and inactive burrows. All soil samples were collected at a depth of approximately 10 cm.

**Fig. 1.**
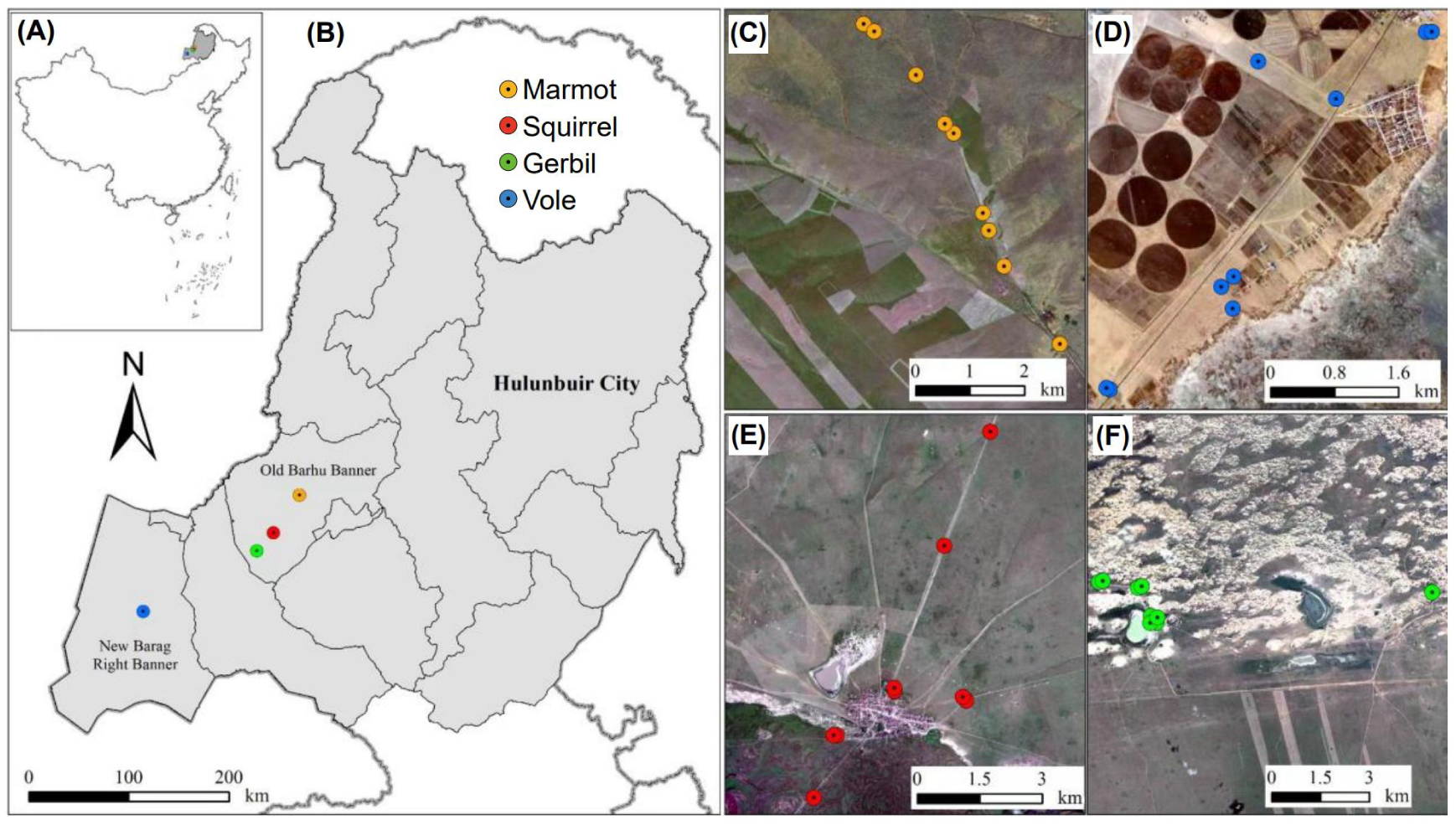
Distribution of sampling sites. (A and B) Four natural rodent habitats in Hulunbuir, China were selected as sampling locations. Nine sampling sites were selected in each rodent habitat of (C) marmot, (D) vole, (E) squirrel, and (F) gerbil, some of which overlapped slightly in the photos due to their proximity. Each sampling site includes three soil types, each with three replicates, as shown in Fig. S1.

At each soil sampling point, a 10 m × 10 m plot was established. Three 1 m × 1 m quadrats were arranged along the diagonal of the plot. Within each quadrat, plant species composition was recorded, and the abundance of each species was quantified by counting or estimating the number of individuals. Plant abundances were classified into seven levels following the grading criteria of Braun-Blanquet (Table S1).

Each replicate soil sample was homogenized and divided into three subsamples for different analyses. For DNA extraction, a 10 g subsample was transported on dry ice and stored at −80 °C as soon as possible. For metal concentration analysis, a 200 g subsample was transported on ice and stored at 4 °C. For soil chemical property analysis, a 100 g subsample was also transported on ice and stored at 4 °C.

Plasma Mass Spectrometry and Atomic Fluorescence Spectroscopy were used to determine the concentrations of Iron (Fe), Calcium (Ca), Magnesium (Mg), Zinc (Zn), Copper (Cu), Manganese (Mn), Nickel (Ni), Cobalt (Co), Chromium (Cr), Titanium (Ti), Vanadium (V), Selenium (Se), Molybdenum (Mo) (Ammann, 2007; Sanchez-Rodas et al., 2010). Soil water contents were measured by oven-drying soil samples at 105 °C for 24 h. Soil pH and electrical conductivity were determined using a pH meter (PHS-2F, INESA Scientific Instrument Co., Ltd., Shanghai, China) and a conductivity meter (DDS-11A, INESA Scientific Instrument Co., Ltd., Shanghai, China), respectively. The potassium dichromate oxidation method was used to measure soil organic matter (Walinga et al., 1992). The concentrations of nitrate (NO ^−^-N), ammonium, available potassium, and available phosphorus were determined according to standardized colorimetry methods (Willey et al., 1999).

### 2.2. DNA extraction, quantification, and sequencing

Total soil DNA was extracted from soil samples using E.Z.N.A.® Soil DNA Kit (Omega Bio-tek, Norcross, GA, U.S.) according to manufacturer’s protocols. The copy numbers of 16S rRNA gene and 18S rRNA gene were quantified by quantitative real-time polymerase chain reaction (qPCR) with at least three technical replicates per sample. Microbial relative and absolute abundances were determined using high-throughput absolute quantification sequencing. The primers 341F (5’-CCTAYGGGRBGCASCAG-3’) and 806R (5’-GGACTACNNGGGTATCTAAT-3’) were used to target the V3-V4 region of bacterial 16S rRNA gene. The primers TAReuk454FWD1 (5’-CCAGCASCYGCGGTAATTCC-3’) and TAReukREV3 (5’-ACTTTCGTTCTTGATYRA-3’), suitable for profiling amoeba diversity, were used to target the V4 region of 18S rRNA gene (Stoeck et al., 2010; Zheng et al., 2022). Amplification conditions for 16S rRNA gene comprised 95 °C for 5 min, followed by 28 cycles of 95 °C for 30 s, 57 °C for 30 s, and 72 °C for 45 s, and a final extension at 72 °C for 10 min. To maximize amplification coverage and specificity to the V4 region of eukaryotes, amplification for 18S rRNA gene was performed twice: once with an annealing temperature of 55 °C for 35 cycles and once at 47 °C for 33 cycles. All other amplification conditions were the same as those used for 16S rRNA gene. The amplification products of 18S rRNA gene from the two reactions were combined and used for sequencing library construction. High-throughput DNA sequencing was performed on an Illumina platform (Shanghai BIOZERON Co., Ltd., Shanghai, China).

### 2.3. Data processing and analysis

Raw reads were qualified using QIIME2 and clustered into operational taxonomic units (OTUs) using UPARSE (version 7.1, http://drive5.com/uparse/) with a 97 % sequence similarity cutoff. The relative abundances of OTUs were transformed to the absolute abundances by multiplying the gene copies per gram soil in each sample. The taxonomy of each 16S rRNA gene sequence was analyzed using RDP Classifier (http://rdp.cme.msu.edu/) based on the silva 16S rRNA gene database (SSU138.1) with a confidence threshold of 80% (Amato et al., 2013). The phylogeny affiliation of 18S rRNA gene was obtained by alignment against the Nucleotide database (version 202003 and 202307). The rooted phylogenetic tree of OTU representative sequences was constructed using QIIME2. Faith’s phylogenetic diversity (PD) of microbial community was calculated using R package *picante*. Multiple comparisons between samples were tested using the Tukey’s HSD method in R package *agricolae* for rRNA gene copies, PD, and soil factors.

A random forest model was constructed to predict important soil factors and OTUs contributing to between-group difference and was validated by 10-fold cross-validation with three replicates using R packaged *MegaR* (Dhungel et al., 2021). The contribution of specific soil properties that showed significant between-group differences on microbial communities were further analyzed using redundancy analysis (RDA) (https://github.com/vegandevs/vegan). The multi-collinear soil factors were removed before RDA analysis based on variance inflation factor less than 10. The significance of the environmental effects on microbial communities in RDA analysis was tested using permutation test. An indicator value was calculated to represent the association strength between each important OTU identified through the random forest model and a particular group or combinations of any two groups using R package *indicspecies*. Differences in microbial communities among groups were assessed via the analysis of similarities (ANOSIM).

Important indicator OTUs were used to construct co-occurrence networks using the R package Weighted Correlation Network Analysis (WGCNA). The co-occurrence network between 16S and 18S OTUs were separately constructed using the indicator OTUs specific to active burrow, inactive burrow, and off-burrow soils. The similarity matrix of microbial community based on Pearson correlation coefficients was transformed into an adjacency matrix using a soft threshold, which was then used to construct a weighted signed co-occurrence network. The filtering criterion for the soft threshold is to ensure that the scale-free network topology structure has an *R*^2^ value greater than 0.85, and the soft threshold value is less than 30. The 16S indicator OTUs strongly co-occurring with amoebae were identified based on their association strength with amoebae OTUs greater than 0.2 for the indicator OTUs of active burrow, inactive burrow, and off-burrow soils.

Bacterial functional profiles were predicted using the R package Tax4Fun2 based on 16S rRNA gene sequences and the KEGG Orthology (KO) database. OTU representative sequences were matched to reference genomes in the NCBI RefSeq database using BLAST. Functional profiles were inferred from the closest matches, and OTU abundances were normalized by the genomic 16S rRNA gene copy number of each microbial taxon. Predicted functions were further summarized at KEGG pathway levels (Wemheuer et al., 2020). Kruskal-Wallis rank sum test was performed to analyze the statistical differences of functional terms. Correlation and clustering relationships between pairs of bacteria and amoebae were displayed using R package *pheatmap*, and co-occurring bacterial and amoebal OTUs were merged when their phylogenetic affiliations were consistent.

The differences in plant communities across different samples were analyzed via Principal Coordinates Analysis (PCoA) using R package *vegan*, and visualized using R package *scatterplot3d*. A distance-based ordination approach was used to reduce each variable set (spatial, soil, and plant variables, as well as absolute abundances and predicted functional abundances of 16S OTUs co-occurring with amoebae) into a one-dimensional ecological feature. Briefly, Bray-Curtis dissimilarities were calculated for each variable based on observed values, followed by PCoA. The first three PCoA axes were weighted by their respective eigenvalues and summed to generate a single composite score for each variable. Subsequently, the one-dimensional ecological features were used for both structural equation modeling (SEM) and mediation analyses, which were conducted using IBM SPSS AMOS (version 21; IBM SPSS, Somers, USA) and the R package *mediation*, respectively. The Pearson correlations between the ecological features and the observed values of the corresponding variables were calculated using R package *vegan*, with significance assessed by permutation tests. The distribution patterns of the ecological features and the observed values of variables significantly correlated (p < 0.001) with the corresponding ecological features were visualized using R package *pheatmap*.

## 3. Results

### 3.1. Soil NO ^−^-N enrichment in active burrows is associated with reduced microbial PD

Microbial abundances, as indicated by 16S and 18S rRNA gene copy numbers, did not differ significantly among active burrow, inactive burrow, and off-burrow soils (Fig. 2A). In contrast, active burrow soils exhibited significantly lower prokaryotic and eukaryotic PD than off-burrow soils, with inactive burrow soils showing intermediate PD (Fig. 2B). Active burrow soils also had significantly higher NO_3_^−^-N and available phosphorus levels compared with both inactive burrow and off-burrow soils, and lower organic matter content than off-burrow soils (Fig. 2C). Other soil physicochemical properties and metal concentrations did not differ significantly across the three soil types (Fig. S2). Random forest analysis identified NO_3_^−^-N as the most important predictor of soil types (Figs. 2D-E). Redundancy analysis indicated that soil organic matter and NO_3_^−^-N each explained a substantial fraction of the variance in both prokaryotic and eukaryotic communities, with NO ^−^-N positively associated with the variation in microbial community composition in specific active burrow samples (Fig. 2F, Table S2).

**Fig. 2.**
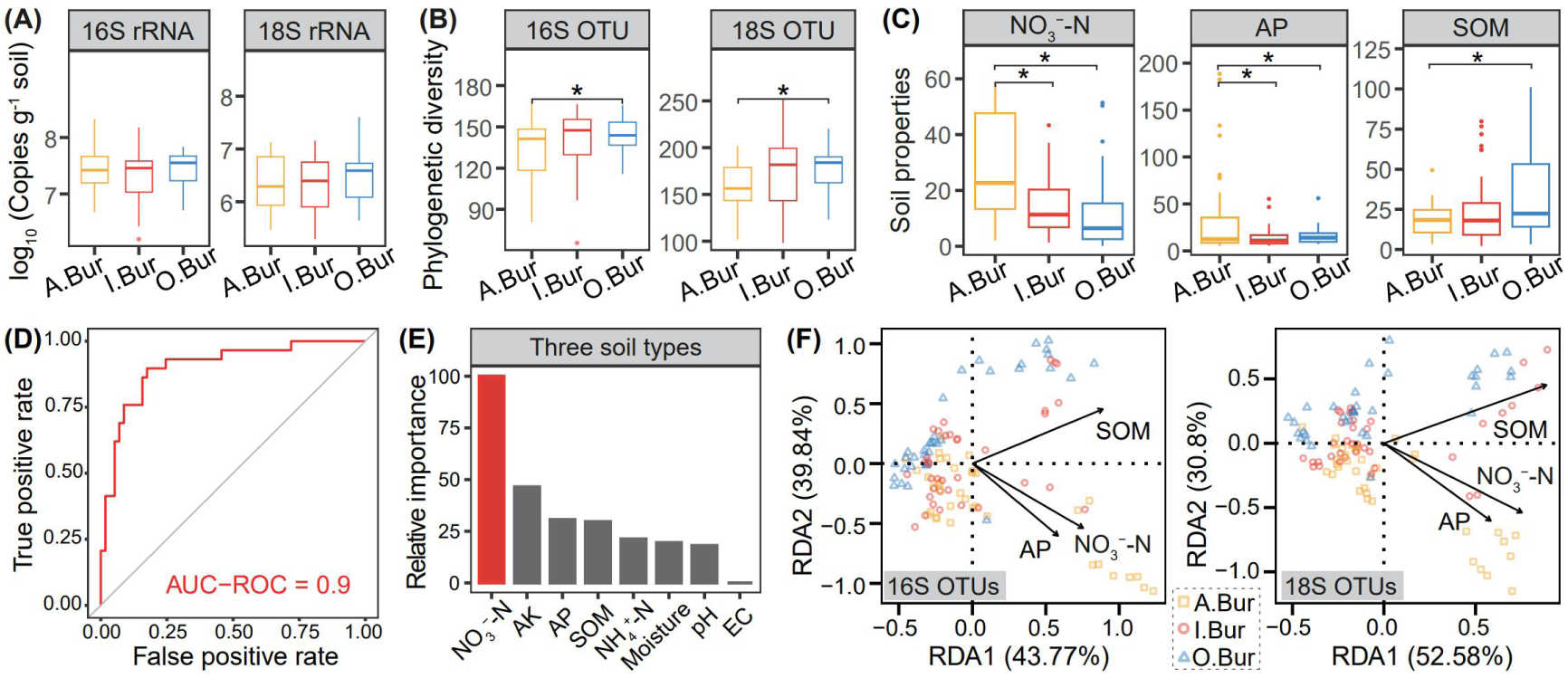
Links between microbial diversity and soil factors in active burrow soils (A.Bur), inactive burrow soils (I.Bur), and off-burrow soils (O.Bur). Differences in (A) microbial absolute abundances, (B) phylogenetic diversity, and (C) soil factors among the three soil types. (D and E) A random forest model identifies the relative importance of soil properties to soil differences among soil types. The predictive performance of the random forest model was evaluated using the area under the receiver operating characteristic curve (AUC-ROC). The colored bar indicates the most important soil factor. (F) Redundancy analysis (RDA) identifies the influences of soil properties on microbial communities. NO_3_^−^-N, nitrate. AK, available potassium. AP, available phosphorus. SOM, soil organic matter. NH_4_^+^-N, ammonium. EC, electrical conductivity. The measurement unit for NO_3_^−^-N and AP is mg‧kg^-1^. The measurement unit for SOM is g‧kg^-1^. Significance level in Figs. 2B-C: 0.05, *.

The effects of rodent species on microbial communities varied among the three soil types (Figs. S3A-B, Table S3). Specifically, marmot active burrow soils showed significantly higher microbial abundance compared to off-burrow, and lower PD compared to inactive burrow soils. In squirrel active burrow soils, the abundance and PD of eukaryotic microbes were significantly lower than that in inactive burrow soils. Gerbil inactive burrow soils exhibited significantly lower microbial abundance and PD relative to off-burrow soils. For voles, microbial abundance showed a significant increasing trend from active burrow to off-burrow soils. In addition, soil NO ^−^-N content exhibited a significant decreasing trend from active burrow soils to off-burrow soils across all four rodent species, whereas patterns of available phosphorus and organic matter varied among species (Fig. S3C, Table S3). Moreover, the positive linear relationship between soil NO_3_^−^-N content and microbial abundance, as well as the negative relationship between NO_3_^−^-N and PD, was significant only in marmot samples (Figs. S3D-E, Table S4).

Overall, active burrow soils exhibited distinct chemical properties and reduced microbial PD compared with off-burrow soils. NO_3_^−^-N was identified as a key predictor of soil type and microbial community variation. However, significant relationships between NO_3_^−^-N and microbial PD were observed only in marmot samples, with other rodent species showing differing patterns.

### 3.2. Taxonomic composition and predicted functions of amoeba-co-occurring indicator bacterial assemblages

To examine microbial community differentiation associated with rodent activity, we applied random forest and indicator species analyses to identify indicator taxa for active burrow, inactive burrow, and off-burrow soils. At the OTU level, the strongest and most consistent differences in both prokaryotic and eukaryotic indicator assemblages were observed between active burrow and off-burrow soils across all rodent species (Table S5).

Differences in indicator assemblages between inactive burrows and off-burrow soils were also significant but generally weaker than those between active burrow and off-burrow soils. In contrast, comparisons between active and inactive burrows were predominantly non-significant for prokaryotic assemblages, whereas eukaryotic assemblages remained significant, albeit with variable effect sizes as measured by ANOSIM *R*.

Given the close ecological interactions between bacteria and amoebae in soils, subsequent analyses were focused on co-occurring bacterial and amoebal taxa within the indicator assemblages associated with each soil type. Inactive burrow indicator OTUs (Ind_I) exhibited a comparatively broader representation of co-occurring bacterial phyla and amoebal genera relative to active burrow indicator OTUs (Ind_A) and off-burrow indicator OTUs (Ind_O) (Figs. S4A-B). At the phylum level, amoeba-co-occurring bacterial assemblages in both Ind_A and Ind_I differed significantly from those in Ind_O, but did not differ significantly from each other (Table S6). These results suggest that burrow environments shape distinct bacterial assemblages while maintaining a core set of indicator bacterial taxa common to both active and inactive burrows.

To assess the functional implications of these taxonomic differences in amoeba-co-occurring bacterial assemblages, we analyzed predicted bacterial functions based on 16S rRNA gene data. Functional predictions indicated that the relative abundances of KEGG nitrogen metabolism-related functional pathways (N metabolism) were higher than those of KEGG infectious disease-related functional pathways (IDFP; Figs. S5 and S6). Moreover, amoeba-co-occurring bacterial assemblages in Ind_A and Ind_I showed higher predicted abundances of IDFP in active and inactive burrow samples, whereas assemblages in Ind_O were enriched in N metabolism in off-burrow samples. In line with these patterns, the pathogenic amoeba *Balamuthia mandrillaris* showed positive correlations with bacterial taxa reported to be associated with human or animal diseases in active burrows (Fig. 3; Tables S7 and S8). In off-burrow soils, amoebae were primarily correlated with nitrogen-cycling bacteria, including nitrogen-fixing and nitrite-oxidizing taxa such as *Beijerinckiaceae* and *Nitrospira*.

**Fig. 3.**
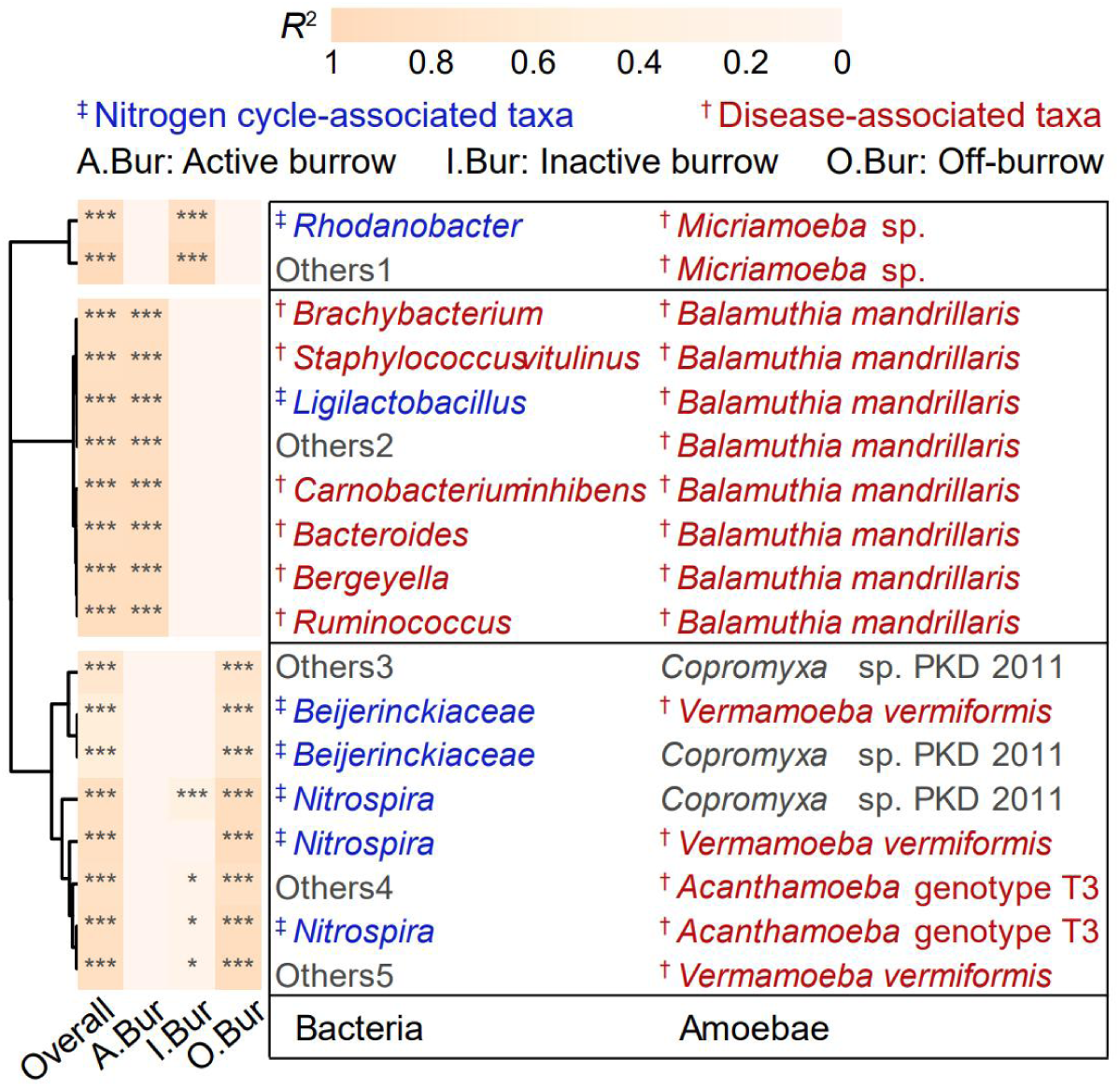
Linear correlation patterns between pairs of co-occurring indicator OTUs specific to soil types with strongest association strengths in the WGCNA networks. Bacteria with uncertain traits were classified as Others which were defined in Table S7. Microbes were colored by functional traits. The references for determining microbial functional traits were listed in Table S8. Disease-associated taxa refer to these taxa associated with human or animal diseases. *R*^2^, adjusted R-squared. *p* < 0.05, *; *p* < 0.01, **; *p* < 0.001, ***.

Taken together, these results indicated that rodent activity influenced both the taxonomic composition and predicted functional profiles of amoeba-co-occurring bacterial assemblages, which exhibited distinct functional signatures: off-burrow soils were enriched in N metabolism, whereas burrow soils retained N metabolism but showed relatively higher representation of predicted infectious disease-related functions.

### 3.3. Interactions among environmental, microbial structural and functional variables

To elucidate the ecological pathways linking rodent activity and environmental factors (soil characteristics, plant composition, and geographical distance), as well as bacterial structure and predicted functions, SEM was applied using one-dimensional ecological features. Results revealed that geographic distance among sampling sites indirectly influenced plant community variation via soil physicochemical properties and metal concentrations (Fig. 4A, Fig. S7). However, rodent species-associated differentiation of plant communities was reduced in burrow habitats, as reflected by lower ANOSIM *R* values compared with off-burrow soils (Fig. S8). Consistent with earlier observations of convergence in amoeba-co-occurring bacterial composition in burrow soils (Table S6), these patterns suggest that rodent-driven soil modification attenuates broader spatial and edaphic controls on both above- and belowground biotic communities.

**Fig. 4.**
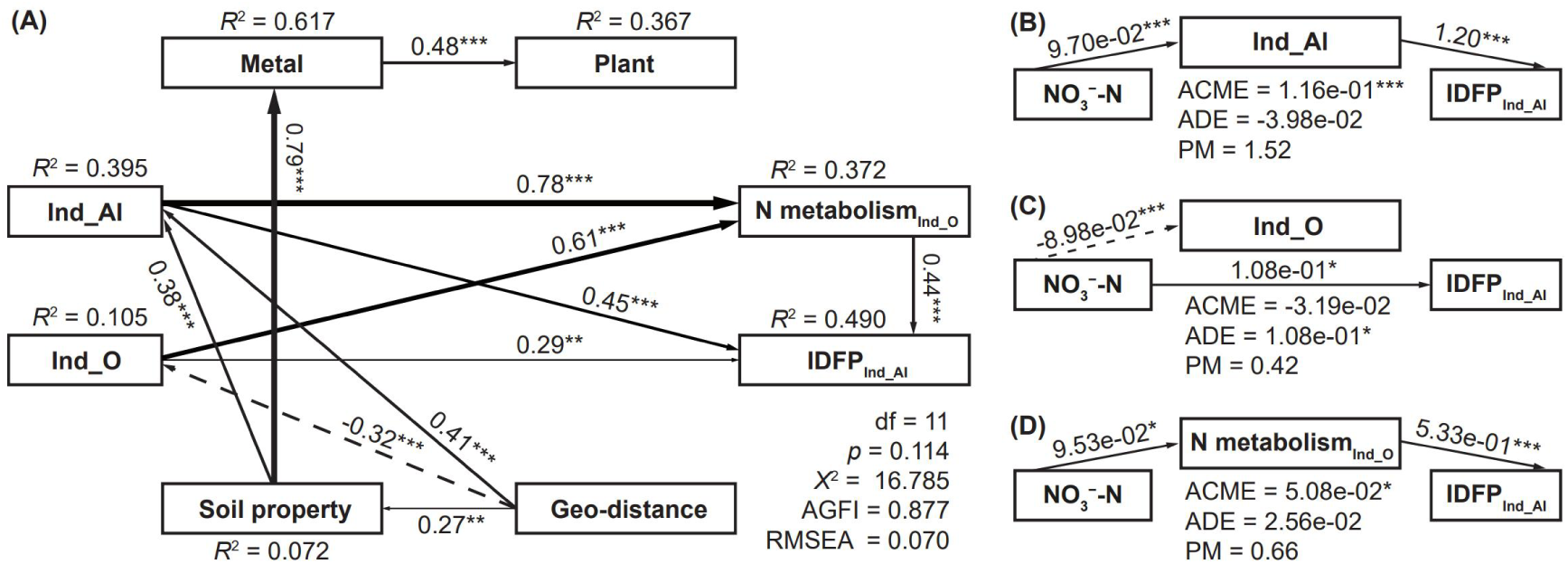
Individual and interactive effects of environmental and microbial factors on microbial functional profiles. (A) A structural equation model and (B-D) mediation analysis depict relationships among multiple variables using their one-dimensional ecological features. Standardized direct effects in structural equation model, direct effects in mediation analysis, and the significance level of direct effects were displayed next to the arrows. *R*^2^, squared multiple correlations. df, degree of freedom. *X*^2^, chi-square. *p* < 0.05, *; *p* < 0.01, **; *p* < 0.001, ***. AGFI, adjusted goodness-of-fit. RMSEA, root mean square error of approximation. ACME, average causal mediation effect. ADE, average direct effect. PM, proportion mediated. Ind_AI, amoeba-co-occurring bacteria identified as burrow indicator OTUs. Ind_O, amoeba-co-occurring bacteria identified as off-burrow indicator OTUs. N metabolism, nitrogen metabolism-related functional pathways. IDFP, infectious diseases-associated functional profiles.

Geographic distance was negatively associated with amoeba-co-occurring bacterial Ind_O. By contrast, both geographic distance and soil properties were positively associated with amoeba-co-occurring bacterial Ind_AI (OTUs identified as Ind_A or Ind_I; Fig. 4A). In turn, bacterial assemblage-level variations were positively associated with N metabolism of amoeba-co-occurring bacterial Ind_O (N metabolism_Ind_O_) and IDFP of amoeba-co-occurring bacterial Ind_AI (IDFP_Ind_AI_). Standardized total effects further indicated that soil properties exerted stronger influences on both N metabolism_Ind_O_ and IDFP_Ind_AI_ than geographic distance (Fig. S9). Moreover, N metabolism_Ind_O_ explained 44% of the variation in IDFP_Ind_AI_. Overall, these results support a network of associations in which soil properties, the composition of amoeba-co-occurring bacterial Ind_AI and Ind_O, and N metabolism_Ind_O_ are primarily linked to IDFP_Ind_AI_.

Mediation analysis based on one-dimensional ecological features indicated that Ind_AI and N metabolism_Ind_O_ significantly mediated the effect of soil NO_3_^−^-N on IDFP_Ind_AI_ (Figs. 4B-D, Fig. S10). To facilitate biological interpretation of these relationships, we compared ecological features with the observed values of the corresponding variables. The NO ^−^-N content and the predicted abundance of IDFP_Ind_AI_ were higher in burrow than in off-burrow soils, whereas their ecological features exhibited the opposite distribution pattern (Fig. S10). By contrast, the ecological features of N metabolism_Ind_O_, Ind_O, and Ind_AI were generally consistent with their observed values. Accordingly, increasing NO ^−^-N was associated with decreases in Ind_AI and N metabolism_Ind_O_, which in turn were linked to higher IDFP_Ind_AI_. Moreover, the lower abundance of Ind_AI and Ind_O, together with the relatively high values of their associated functional profiles, indicate that variation in predicted functional profiles was not closely associated with bacterial abundance, suggesting potential shifts in the composition of amoeba-associated assemblages across soil environments.

Taken together, these results support a model in which NO ^−^-N enrichment is indirectly associated with increased infectious disease-related functional potential in amoeba-co-occurring bacterial assemblages through changes in assemblage structure and nitrogen cycling-related functional potential, rather than through bacterial abundance.

## 4. Discussion

In this study, we examined soils associated with burrows of four rodent species to understand how rodent activity shapes both soil characteristics and microbial interactions. Our findings demonstrate that while multiple soil properties varied in a rodent species-specific manner, soil NO_3_^−^-N consistently responded to rodent activity across all species, highlighting its central role in shaping microbial community structure. Specifically, rodent activities in burrows elevated NO ^−^-N levels, reduced microbial PD, and altered amoeba–bacteria interactions, regardless of the rodent species (Fig. 5).

**Fig. 5.**
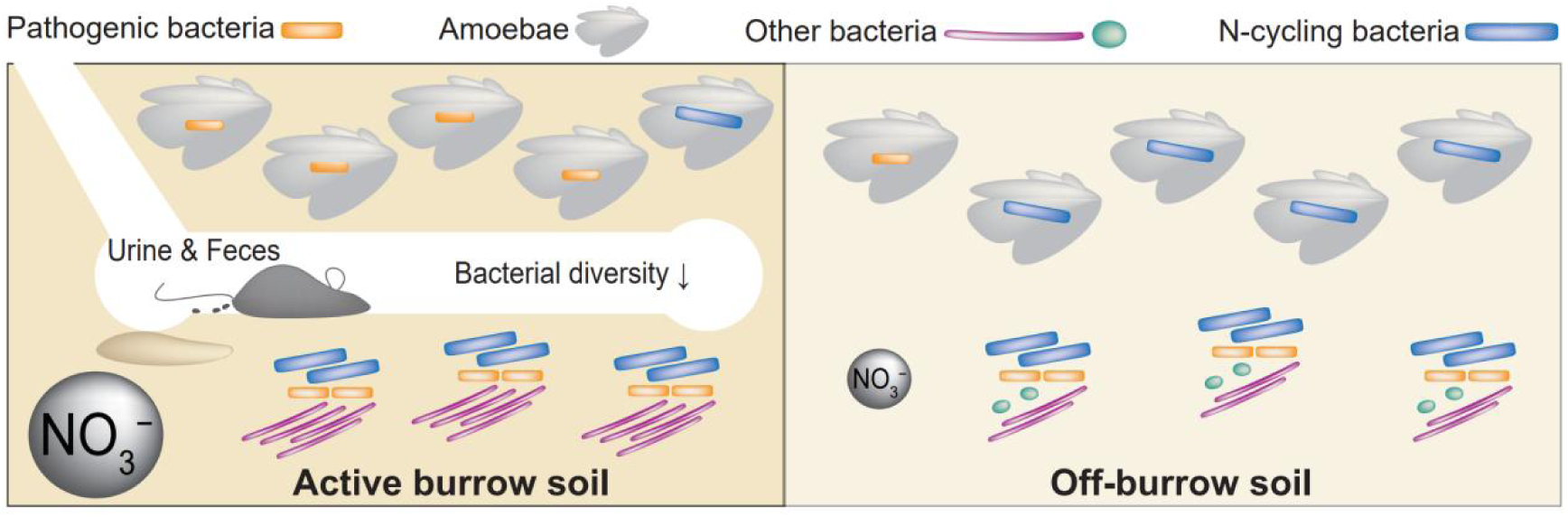
Model illustrating rodent burrows as a potential source of pathogen emergence.

Consistent with previous studies, our data revealed that NO_3_^−^-N content was higher in active burrow than in off-burrow soils across all rodent species (Ambus et al., 2007; Clark et al., 2005; Clark et al., 2016). Rodent foraging and excretion behaviors are known to influence the spatial distribution of soil nitrogen (Clark et al., 2016; Kuznetsova et al., 2013). The accumulation of urine and feces in active burrows likely contributes to the high NO ^−^-N concentration. In contrast, patterns of variation in other measured soil factors among active burrow, inactive burrow, and off-burrow soils differed across rodent species. This species-specificity may reflect differences in the habitats occupied by each rodent species, which are characterized by distinct vegetation types (Kuznetsova et al., 2013; Yoshihara et al., 2010). Although rodent activities tended to homogenize plant communities among the four rodent habitats (Fig. S8), local environmental variation maintained ecosystem differentiation within burrow-associated habitats (Kuypers et al., 2018; Yoshihara et al., 2010). Overall, the effect of rodent activity on soil NO ^−^-N follows consistent cross-species niche differentiation, while its effect on other soil factors exhibits species-specific variations.

Active burrow soils exhibited lower microbial PD and no significant difference in microbial abundance compared to off-burrow soils. Moreover, NO ^−^-N content showed opposite correlations with microbial PD and abundance in marmot samples, indicating that NO_3_^−^-N enrichment may restructure microbial community composition without uniformly reducing microbial biomass. These patterns are consistent with niche-based selection and environmental filtering processes that favor phylogenetically clustered assemblages adapted to nitrogen-enriched conditions (Gao et al., 2025; Horner-Devine and Bohannan, 2006). Beyond abiotic filtering, modern coexistence theory predicts that phylogenetic clustering can arise through biotic filtering when competitively superior lineages exclude distantly related taxa under favorable resource conditions (Goberna et al., 2014). Consistent with this theory, increased nitrogen availability in burrow soils may strengthen the dominance of particular microbial taxa and increase the phylogenetic relatedness of coexisting taxa. In addition, microbial predators such as amoebae can closely interact with soil bacteria under varying environmental conditions, thereby influencing microbial survival and coexistence dynamics (Cecil and Yoder-Himes, 2024; Van Heijnsbergen et al., 2016). Together, these findings suggest that elevated soil NO_3_^−^-N associated with rodent activity may structure microbial assemblages through the combined effects of environmental filtering and microbial interactions.

Further supporting this interpretation, soil NO ^−^-N was also associated with pronounced shifts in amoeba–bacteria co-occurrence patterns. In off-burrow soils, amoebae were typically associated with putative nitrogen-fixing and nitrifying bacteria, in line with previous findings that amoebae can facilitate nitrogen availability by harboring symbiotic bacteria in nitrogen-limited soils (Ekelund and Rønn, 1994; Elliott et al., 1979). In burrow soils characterized by elevated NO_3_^−^-N, amoebae were additionally associated with bacterial taxa linked to IDFP. These associations may reflect shifts in bacterial functional traits favored under high NO ^−^-N conditions, rather than direct selection for pathogenicity. One possible explanation is that some pathogenic bacteria harbor nitrate reductase genes which may enhance their adaptation to local nitrogen conditions through nitrate reduction, a process contributing to nitrogen turnover (Pieper et al., 2008; Zhou et al., 2004). These contrasting association patterns suggest that soil NO_3_^−^-N may shape the selective microenvironment of amoebae, potentially favoring bacterial functional potential associated with pathogenic taxa in rodent burrows. However, these patterns should be interpreted cautiously, as they represent statistical associations based on functional predictions inferred from 16S rRNA data.

Mediation analysis further revealed a potential pathway linking NO_3_^−^-N enrichment to higher IDFP in amoeba-co-occurring bacteria, mediated by shifts in assemblage structure and nitrogen-cycling functional potential. In grasslands, nitrogen supply is predominantly regulated by nitrogen-fixing organisms (Soong et al., 2020). The redistribution of available nitrogen between off-burrow and burrow soils may therefore alter the local nutrient environment and reshape the ecological context in which amoeba–bacteria interactions occur (Cecil and Yoder-Himes, 2024; Clark et al., 2016). Importantly, the shifts in bacterial functional profiles appeared to be decoupled from their abundance, reflecting coordinated changes in community composition and predicted functional attributes. Similar structure–function relationships have been widely documented in soil microbial ecosystems, where shifts in community assembly and interaction networks can strongly influence microbial functional capacities (Zhang et al., 2024; Zhu et al., 2024). These observations support a mediation model in which NO ^−^-N enrichment modulates amoeba-associated bacterial assemblages via indirect ecological pathways rather than through direct effects of bacterial abundance.

Overall, our study indicated that high NO_3_^−^-N levels in rodent burrow soils were associated with shifts in amoeba–bacteria interactions toward bacterial assemblages carrying IDFP, potentially favoring the persistence of taxa with traits relevant to pathogenic lifestyles. However, functional predictions based on 16S data may not fully capture the true functions of the bacterial assemblage, especially when taxa are underrepresented in reference databases or when predicted functions do not reflect *in situ* gene expression. Future studies integrating metagenomics, transcriptomics, and experimental approaches will be necessary to validate these hypotheses and to clarify the ecological and epidemiological relevance of NO_3_^−^-N-driven amoeba–bacteria interactions across diverse ecosystems.

## CRediT authorship contribution statement

**Chunfang Zhang:** Conceptualization, Investigation, Data curation, Formal analysis, Methodology, Visualization, Writing – original draft, Writing – review and editing, Funding acquisition. **Florent Sebbane:** Visualization, Writing – review and editing. **Chutian Zhang:** Investigation, Visualization. **Jason D. Whittington:** Writing – review and editing. **Yong Zhao:** Methodology. **Chaolemen:** Investigation, Resources. **Ruifu Yang:** Methodology, Writing – review and editing. **Nils Christian Stenseth:** Supervision, Writing – review and editing, Funding acquisition. **Lei Xu:** Project administration, Resources, Writing – review and editing, Funding acquisition.

## Supporting information

Supplementary Interactive Plot Data

## Acknowledgements

We thank Yang Wang, Ruier Song, Yuhui Wu, Narisu, Duxi, Xiaoxiang Li, Zhenqi Li, and Hanwei Liang for their expertise and support in the sampling work. We appreciate Zeyuan Pei’s assistance in mediation analysis. This study received support from the National Natural Science Foundation of China (U22A20363), the National Key R&D Program of China (2024YFC2310101). We also acknowledge the support from the Research Fund of Vanke School of Public Health, Tsinghua University. Additional support was provided by the Scientific Research Startup Fund of Zhengzhou University (32213169) and the China Scholarship Council. This work was partially supported by the ERC Synergy project – Synergy-Plague (ERC-2023-SyG, GA no.101118880).

## Appendix A. Supplementary data

Supplementary data to this article can be found online.

## Data availability

The raw 16S and 18S rRNA gene amplicon sequencing data have been deposited in the Genome Sequence Archive (GSA) under BioProject accession number PRJCA037203. Absolute quantitative copy numbers of 16S and 18S rRNA genes were provided as Supplementary Interactive Plot Data (Gene copies.csv).

