## Supplementary material for "Rodent-driven NO_3_^−^-N enrichment reshapes amoeba–bacteria co-occurrence and bacterial functional potential in burrow soils"

**Contents**

**1. Figures**

**Fig. S1** Schematic diagram of soil sampling from each natural habitat of marmot, squirrel, gerbil, and vole (not to scale).

**Fig. S2** Differences in (A) soil properties and (B) metal concentrations among active burrow soils (A.Bur), inactive burrow soils (I.Bur), and off-burrow soils (O.Bur). AK (available potassium) and $NH_4^+$-N (ammonium) are expressed in mg·kg$^{-1}$, electrical conductivity (EC) in μS·cm$^{-1}$, and moisture content as percentage. $Fe_2O_3$, MgO, and CaO are reported as mass percentages of oven-dried soil, and other metal concentrations are expressed in μg·g$^{-1}$. No significant differences were detected among soil types ($p > 0.05$).

**Fig. S3** Variation in microbial communities and their correlations with soil nitrate. Mean values of (A) microbial absolute abundances, (B) phylogenetic diversity, and (C) key soil properties in active burrow soils (A.Bur), inactive burrow soils (I.Bur), and off-burrow soils (O.Bur). Error bars indicate standard error of the mean. Nitrate ($NO_3^-$-N) and available phosphorus (AP) are expressed in mg·kg$^{-1}$, and soil organic matter (SOM) in g·kg$^{-1}$. Linear correlations (D) between $NO_3^-$-N content and microbial absolute abundance, as well as (E) between $NO_3^-$-N content and microbial phylogenetic diversity in marmot samples. $p < 0.05$, *; $p < 0.01$, **; $p < 0.001$, ***.

**Fig. S4** Abundance distribution of co-occurring indicator taxa in different soil types. (A) Phylum-level compositions of indicator bacteria co-occurring with amoebae and (B) genus-level compositions of indicator amoebae co-occurring with bacteria. Relative abundance represents the proportion of each taxon within a given sample group (A.Bur, I.Bur, or O.Bur), calculated separately for each group of indicator OTUs (Ind_A, Ind_I, or Ind_O).

**Fig. S5** Potential disease-associated functional pathways of indicator bacteria co-occurring with amoebae. (A-C) KEGG Orthology (KO) pathway abundances predicted from amoeba-co-occurring bacteria identified as active burrow indicator OTUs (Ind_A), inactive burrow indicator OTUs (Ind_I), and off-burrow indicator OTUs (Ind_O). The bar plots display the relative abundances of bacterial functional profiles in each sample group. Bacterial, P, Viral represent bacterial, parasitic, viral infectious diseases, respectively. $p < 0.05$, *; $p < 0.01$, **; $p < 0.001$, ***.

**Fig. S6** Potential functional pathways associated with nitrogen metabolism of indicator bacteria co-occurring with amoebae. (A-C) KEGG Orthology (KO) pathway abundances predicted from amoeba-co-occurring bacteria identified as active burrow indicator OTUs (Ind_A), inactive burrow indicator OTUs (Ind_I), and off-burrow indicator OTUs (Ind_O). The bar plots display the relative abundances of bacterial functional profiles in each sample group. Bacterial, P, Viral represent bacterial, parasitic, viral infectious diseases, respectively. $p < 0.001$, ***.

**Fig. S7** Complete structural equation model with standardized estimates (A) and unstandardized estimates (B). e1-e7, error variables. AI_eig, one-dimensional ecological feature of the abundance of amoebae-co-occurring bacteria identified as burrow indicator OTUs. O_eig, one-dimensional ecological feature of the abundance of amoebae-co-occurring bacteria identified as off-burrow indicator OTUs. Ncycle_eig, one-dimensional ecological feature of the functional abundances of amoebae-co-occurring bacteria identified as off-burrow indicator OTUs. Disease_eig, one-dimensional ecological feature of the functional abundances of amoebae-co-occurring bacteria identified as burrow indicator OTUs. Soil, soil physicochemical properties. Geodist, the geographical distances between sampling sites. $R^2$, squared multiple correlations. df, degree of freedom. $X^2$, chi-square. AGFI, adjusted goodness-of-fit. RMSEA, root mean square error of approximation.

**Fig. S8** Three-dimensional PCoA (Principal Coordinates Analysis) illustrates the differences in plant communities across four rodent habitats within each soil type. The magnitude of between-group differences was assessed using ANOSIM statistic *R* value. $p < 0.001$, ***.

**Fig. S9** Standardized total effects among environmental factors, microbial factors, and microbial functional profiles. Standardized total effects are the sum of standardized direct effects and indirect effects. Ind_AI, amoeba-co-occurring bacteria identified as burrow indicator OTUs. Ind_O, amoeba-co-occurring bacteria identified as off-burrow indicator OTUs. N metabolism, nitrogen metabolism-related functional pathways. IDFP, infectious diseases-associated functional profiles.

**Fig. S10** A simplified diagram of the distribution patterns of one-dimensional ecological features and their significantly correlated ($p < 0.001$) variable values. The heatmap shows the Zscores of the means of the one-dimensional ecological feature and variable values in burrow and off-burrow samples. Zscore is obtained by subtracting the mean of the values in each column and then dividing it by their standard deviation. $NO_3^-$-N, the one-dimensional ecological feature of soil nitrate contents. $IDFP_{Ind\_AI}$, the one-dimensional ecological feature of the functional profiles associated with infectious diseases of amoeba-co-occurring bacteria identified as burrow indicator OTUs. N $metabolism_{Ind\_O}$, the one-dimensional ecological feature of the functional profiles associated with nitrogen metabolism of amoeba-co-occurring bacteria identified as off-burrow indicator OTUs. Ind_O, the one-dimensional ecological feature of the absolute abundances of amoeba-co-occurring bacteria identified as off-burrow indicator OTUs. Ind_AI, the one-dimensional ecological feature of the absolute abundances of amoeba-co-occurring bacteria identified as burrow indicator OTUs.

### 2. Tables

**Table S1** Grading criteria for plant abundance based on individual counts in 1 m × 1 m quadrats.

**Table S2** Permutation test between soil properties and microbial communities in redundancy analysis (RDA) among active burrow, inactive burrow, and off-burrow soils.

**Table S3** Significance test of the differences of microbial absolute abundances, phylogenetic diversity, and key soil properties among three soil types for each rodent species.

**Table S4** Linear correlation between soil $NO_3^-$-N content and both microbial absolute abundance and phylogenetic diversity in each rodent habitat.

**Table S5** ANOSIM analysis of the between-group differences of important indicator OTUs of three soil types.

**Table S6** ANOSIM analysis of the between-group abundance differences of important indicator bacterial OTUs co-occurring with amoebae at phylum level.

**Table S7** Definitions of the abbreviations in Figure 3.

**Table S8** References for determining microbial functional traits in Figure 3.

**Figure S1**

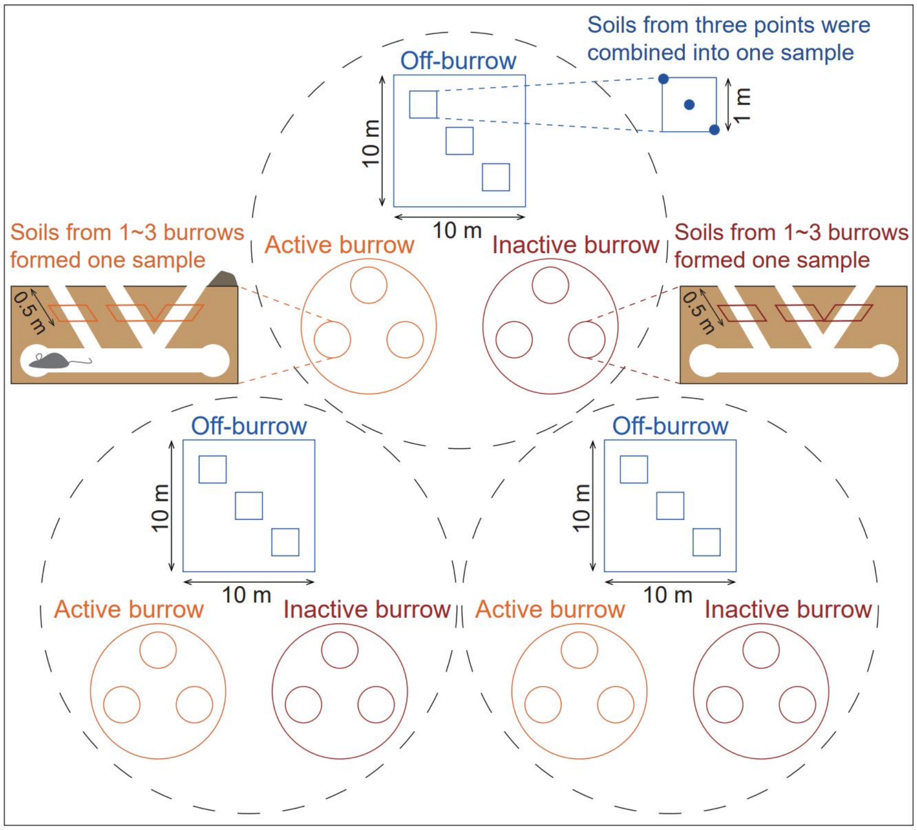


**Fig. S1** Schematic diagram of soil sampling from each natural habitat of marmot, squirrel, gerbil, and vole (not to scale).

**Figure S2**

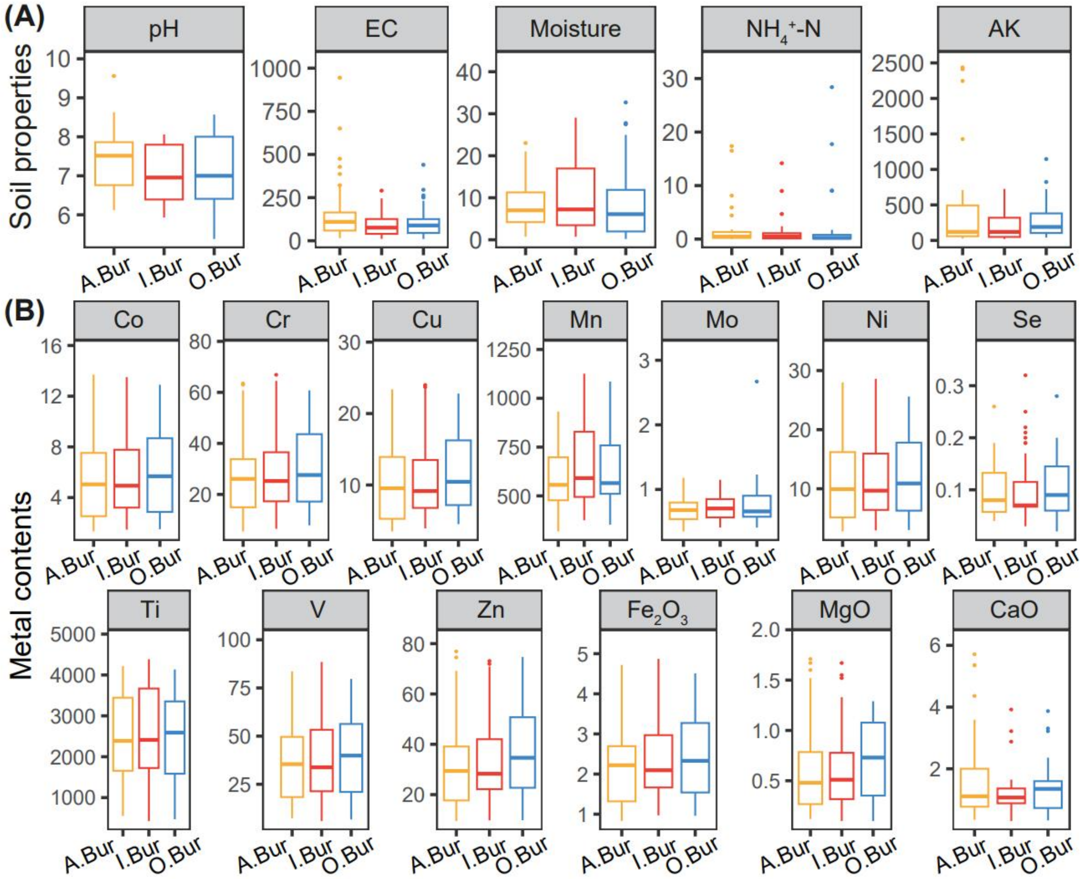


**Fig. S2** Differences in (A) soil properties and (B) metal concentrations among active burrow soils (A.Bur), inactive burrow soils (I.Bur), and off-burrow soils (O.Bur). AK (available potassium) and $NH_4^+$-N (ammonium) are expressed in mg·kg$^{-1}$, electrical conductivity (EC) in µS·cm$^{-1}$, and moisture content as percentage. $Fe_2O_3$, MgO, and CaO are reported as mass percentages of oven-dried soil, and other metal concentrations are expressed in µg·g$^{-1}$. No significant differences were detected among soil types ($p > 0.05$).

### Figure S3

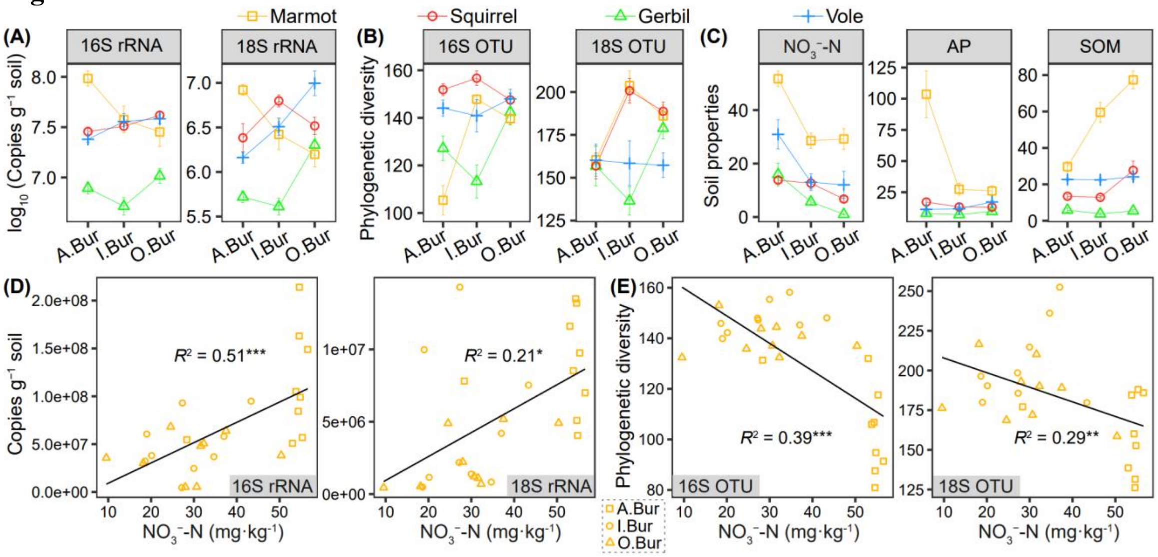


**Fig. S3** Variation in microbial communities and their correlations with soil nitrate. Mean values of (A) microbial absolute abundances, (B) phylogenetic diversity, and (C) key soil properties in active burrow soils (A.Bur), inactive burrow soils (I.Bur), and off-burrow soils (O.Bur). Error bars indicate standard error of the mean. Nitrate ($NO_3^-$-N) and available phosphorus (AP) are expressed in mg·kg$^{-1}$, and soil organic matter (SOM) in g·kg$^{-1}$. Linear correlations (D) between $NO_3^-$-N content and microbial absolute abundance, as well as (E) between $NO_3^-$-N content and microbial phylogenetic diversity in marmot samples. $p < 0.05$, *; $p < 0.01$, **; $p < 0.001$, ***.

**Figure S4**

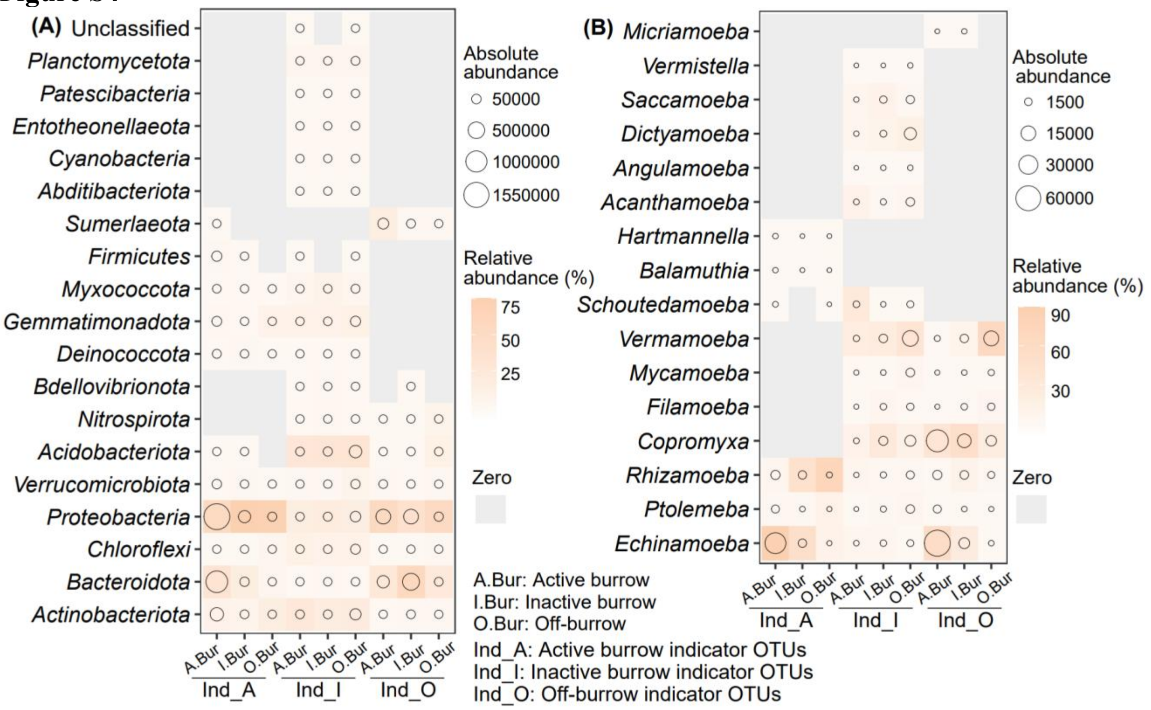


**Fig. S4** Abundance distribution of co-occurring indicator taxa in different soil types. (A) Phylum-level compositions of indicator bacteria co-occurring with amoebae and (B) genus-level compositions of indicator amoebae co-occurring with bacteria. Relative abundance represents the proportion of each taxon within a given sample group (A.Bur, I.Bur, or O.Bur), calculated separately for each group of indicator OTUs (Ind_A, Ind_I, or Ind_O).

**Figure S5**

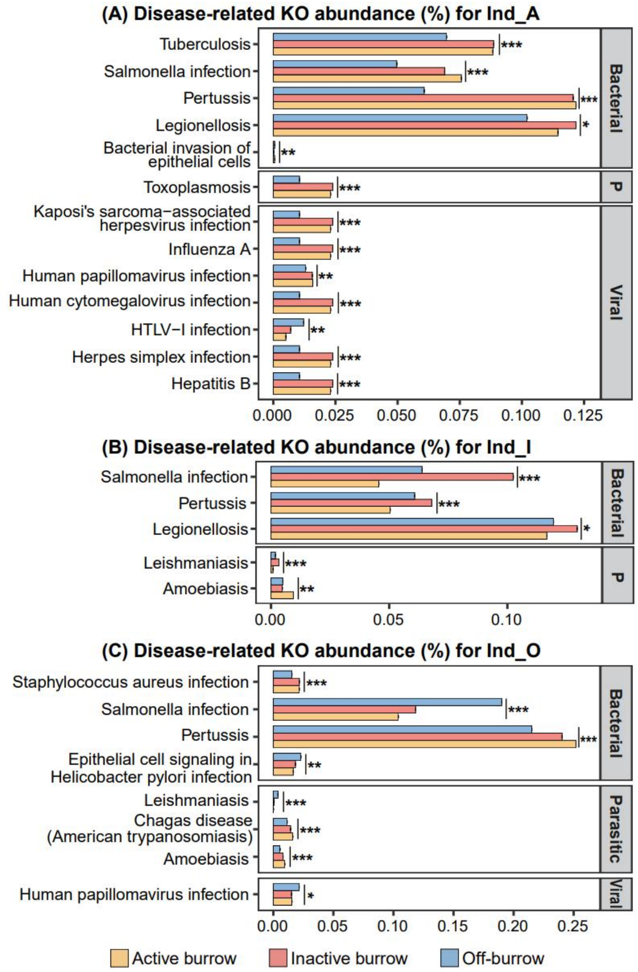


**Fig. S5** Potential disease-associated functional pathways of indicator bacteria co-occurring with amoebae. (A-C) KEGG Orthology (KO) pathway abundances predicted from amoeba-co-occurring bacteria identified as active burrow indicator OTUs (Ind_A), inactive burrow indicator OTUs (Ind_I), and off-burrow indicator OTUs (Ind_O). The bar plots display the relative abundances of bacterial functional profiles in each sample group. Bacterial, P, Viral represent bacterial, parasitic, viral infectious diseases, respectively. $p < 0.05$, *; $p < 0.01$, **; $p < 0.001$, ***.

**Figure S6**

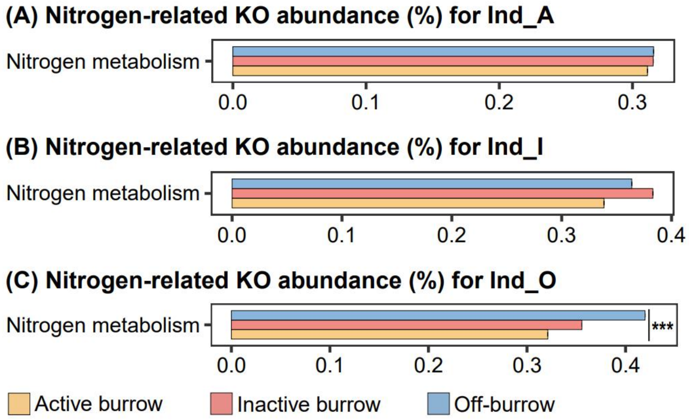


**Fig. S6** Potential functional pathways associated with nitrogen metabolism of indicator bacteria co-occurring with amoebae. (A-C) KEGG Orthology (KO) pathway abundances predicted from amoeba-co-occurring bacteria identified as active burrow indicator OTUs (Ind_A), inactive burrow indicator OTUs (Ind_I), and off-burrow indicator OTUs (Ind_O). The bar plots display the relative abundances of bacterial functional profiles in each sample group. Bacterial, P, Viral represent bacterial, parasitic, viral infectious diseases, respectively. $p < 0.001$, ***.

**Figure S7**

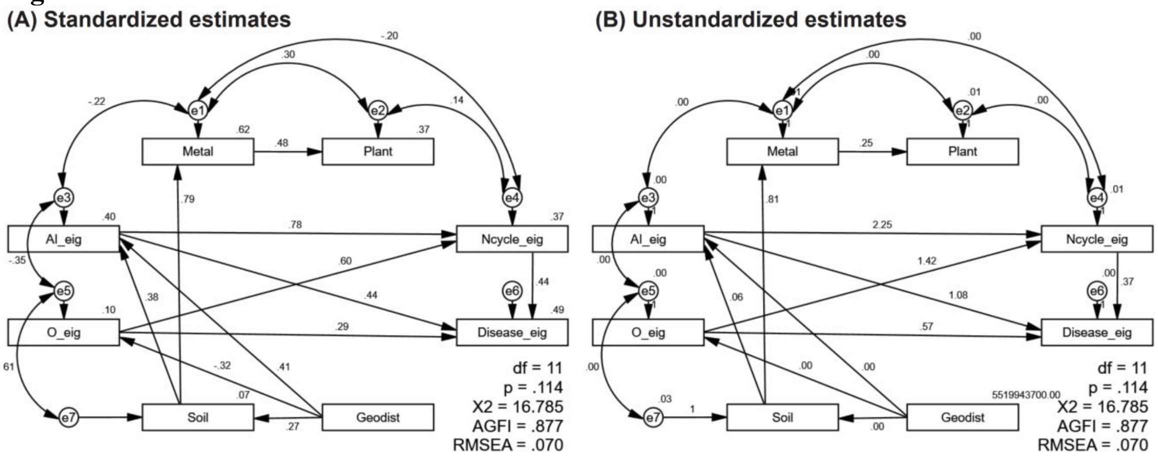


**Fig. S7** Complete structural equation model with standardized estimates (A) and unstandardized estimates (B). e1-e7, error variables. AI_eig, one-dimensional ecological feature of the abundance of amoebae-co-occurring bacteria identified as burrow indicator OTUs. O_eig, one-dimensional ecological feature of the abundance of amoebae-co-occurring bacteria identified as off-burrow indicator OTUs. Ncycle_eig, one-dimensional ecological feature of the functional abundances of amoebae-co-occurring bacteria identified as off-burrow indicator OTUs. Disease_eig, one-dimensional ecological feature of the functional abundances of amoebae-co-occurring bacteria identified as burrow indicator OTUs. Soil, soil physicochemical properties. Geodist, the geographical distances between sampling sites. $R^2$, squared multiple correlations. df, degree of freedom. $X^2$, chi-square. AGFI, adjusted goodness-of-fit. RMSEA, root mean square error of approximation.

**Figure S8**

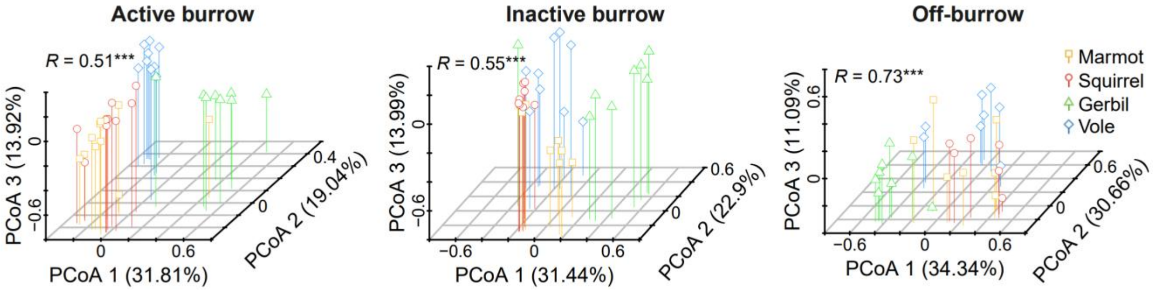


**Fig. S8** Three-dimensional PCoA (Principal Coordinates Analysis) illustrates the differences in plant communities across four rodent habitats within each soil type. The magnitude of between-group differences was assessed using ANOSIM statistic *R* value. $p < 0.001$, ***.

**Figure S9**

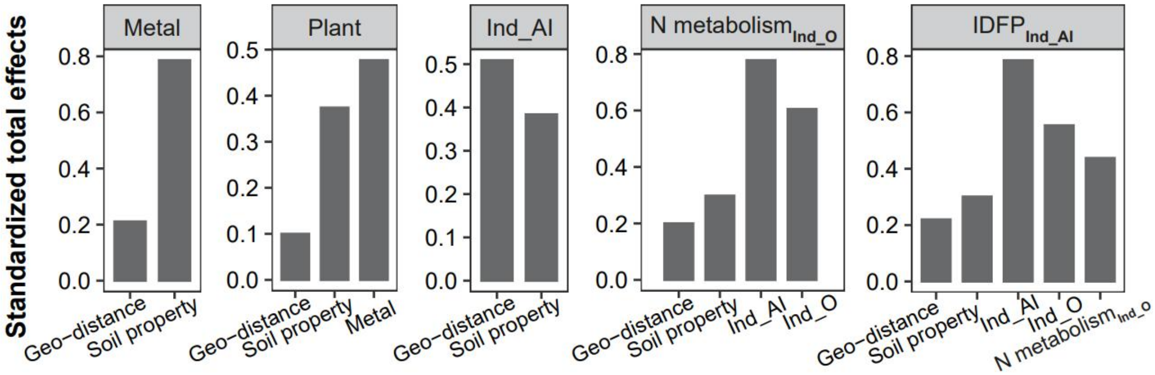


**Fig. S9** Standardized total effects among environmental factors, microbial factors, and microbial functional profiles. Standardized total effects are the sum of standardized direct effects and indirect effects. Ind_AI, amoeba-co-occurring bacteria identified as burrow indicator OTUs. Ind_O, amoeba-co-occurring bacteria identified as off-burrow indicator OTUs. N metabolism, nitrogen metabolism-related functional pathways. IDFP, infectious diseases-associated functional profiles.

**Figure S10**

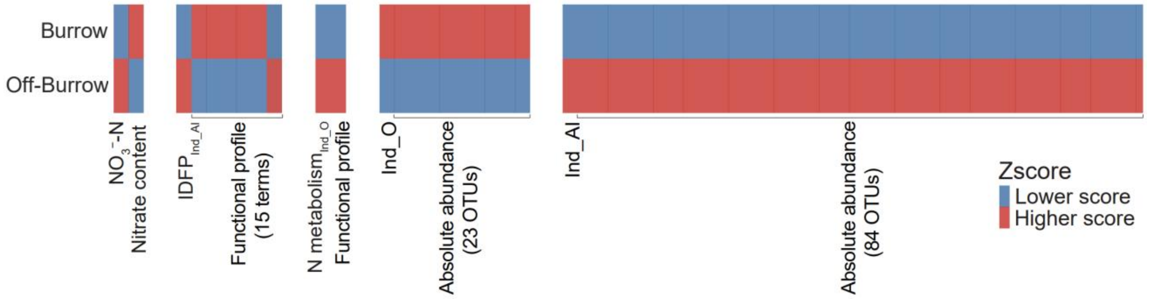


**Fig. S10** A simplified diagram of the distribution patterns of one-dimensional ecological features and their significantly correlated ($p$ < 0.001) variable values. The heatmap shows the Zscores of the means of the one-dimensional ecological feature and variable values in burrow and off-burrow samples. Zscore is obtained by subtracting the mean of the values in each column and then dividing it by their standard deviation. $NO_3^-$-N, the one-dimensional ecological feature of soil nitrate contents. $IDFP_{Ind\_AI}$, the one-dimensional ecological feature of the functional profiles associated with infectious diseases of amoeba-co-occurring bacteria identified as burrow indicator OTUs. N $metabolism_{Ind\_O}$, the one-dimensional ecological feature of the functional profiles associated with nitrogen metabolism of amoeba-co-occurring bacteria identified as off-burrow indicator OTUs. Ind_O, the one-dimensional ecological feature of the absolute abundances of amoeba-co-occurring bacteria identified as off-burrow indicator OTUs. Ind_AI, the one-dimensional ecological feature of the absolute abundances of amoeba-co-occurring bacteria identified as burrow indicator OTUs.

**Table S1** Grading criteria for plant abundance based on individual counts in 1 m × 1 m quadrats.

| **Categories** | **Abundances** | | | | | | |
|---|---|---|---|---|---|---|---|
| Count | 1~2 | 3~4 | 5~7 | 8~15 | 16~30 | 31~50 | 51~100 |
| Level | 1 | 2 | 3 | 4 | 5 | 6 | 7 |

**Table S2** Permutation test between soil properties and microbial communities in redundancy analysis (RDA) among active burrow, inactive burrow, and off-burrow soils.

| **16S OTUs** | | | **18S OTUs** | | |
|---|---|---|---|---|---|
| Soil properties† | $R^2$ | $p$ | Soil properties | $R^2$ | $p$ |
| SOM | 0.85 | <0.001 | SOM | 0.88 | <0.001 |
| $NO_3^-$-N | 0.72 | <0.001 | $NO_3^-$-N | 0.69 | <0.001 |
| AP | 0.62 | <0.001 | AP | 0.56 | <0.001 |

† SOM, soil organic matter; $NO_3^-$-N, nitrate; AK, available potassium.

**Table S3** Significance test of the differences of microbial absolute abundances, phylogenetic diversity, and key soil properties among three soil types for each rodent species.

| **Rodents** | **Comparison groups**[†] | **Gene abundance** | | **Phylogenetic diversity** | | **Soil properties** | | |
|---|---|---|---|---|---|---|---|---|
| | | **16S rRNA** | **18S rRNA** | **16S OTU** | **18S OTU** | **$NO_3^-$-N** | **AP** | **SOM** |
| Marmot | A.Bur vs I.Bur | NS[‡] | * | * | * | * | * | * |
| | A.Bur vs O.Bur | * | * | * | NS | * | * | * |
| | I.Bur vs O.Bur | NS | NS | NS | NS | NS | NS | * |
| Squirrel | A.Bur vs I.Bur | NS | * | NS | * | NS | NS | NS |
| | A.Bur vs O.Bur | * | NS | NS | * | * | NS | * |
| | I.Bur vs O.Bur | NS | NS | NS | NS | * | NS | * |
| Gerbil | A.Bur vs I.Bur | NS | NS | NS | NS | * | NS | NS |
| | A.Bur vs O.Bur | NS | * | NS | NS | * | NS | NS |
| | I.Bur vs O.Bur | * | * | * | * | NS | NS | NS |
| Vole | A.Bur vs I.Bur | NS | NS | NS | NS | * | NS | NS |
| | A.Bur vs O.Bur | * | * | NS | NS | * | * | NS |
| | I.Bur vs O.Bur | NS | * | NS | NS | NS | * | NS |

[†] The A.Bur, I.Bur, and O.Bur represent active burrow soils, inactive burrow soils, and off-burrow soils, respectively.
[‡] NS represents not significant at 0.05.

**Table S4** Linear correlation between soil $NO_3^-$-N content and both microbial absolute abundance and phylogenetic diversity in each rodent habitat.

| **Rodents** | **Independent variable** | **Dependent variable** | $R^2$ | $p$ |
|---|---|---|---|---|
| Marmot | $NO_3^-$-N ($mg \cdot kg^{-1}$) | 16S rRNA gene (Copies $g^{-1}$ soil) | 0.390 | <0.001 |
| | $NO_3^-$-N ($mg \cdot kg^{-1}$) | 18S rRNA gene (Copies $g^{-1}$ soil) | 0.286 | 0.004 |
| | $NO_3^-$-N ($mg \cdot kg^{-1}$) | Phylogenetic diversity (16S OTUs) | 0.510 | <0.001 |
| | $NO_3^-$-N ($mg \cdot kg^{-1}$) | Phylogenetic diversity (18S OTUs) | 0.212 | 0.016 |
| Squirrel | $NO_3^-$-N ($mg \cdot kg^{-1}$) | 16S rRNA gene (Copies $g^{-1}$ soil) | 0.005 | NS† |
| | $NO_3^-$-N ($mg \cdot kg^{-1}$) | 18S rRNA gene (Copies $g^{-1}$ soil) | 0.004 | NS |
| | $NO_3^-$-N ($mg \cdot kg^{-1}$) | Phylogenetic diversity (16S OTUs) | 0.010 | NS |
| | $NO_3^-$-N ($mg \cdot kg^{-1}$) | Phylogenetic diversity (18S OTUs) | 0.005 | NS |
| Gerbil | $NO_3^-$-N ($mg \cdot kg^{-1}$) | 16S rRNA gene (Copies $g^{-1}$ soil) | <0.001 | NS |
| | $NO_3^-$-N ($mg \cdot kg^{-1}$) | 18S rRNA gene (Copies $g^{-1}$ soil) | 0.069 | NS |
| | $NO_3^-$-N ($mg \cdot kg^{-1}$) | Phylogenetic diversity (16S OTUs) | 0.004 | NS |
| | $NO_3^-$-N ($mg \cdot kg^{-1}$) | Phylogenetic diversity (18S OTUs) | 0.004 | NS |
| Vole | $NO_3^-$-N ($mg \cdot kg^{-1}$) | 16S rRNA gene (Copies $g^{-1}$ soil) | 0.029 | NS |
| | $NO_3^-$-N ($mg \cdot kg^{-1}$) | 18S rRNA gene (Copies $g^{-1}$ soil) | 0.036 | NS |
| | $NO_3^-$-N ($mg \cdot kg^{-1}$) | Phylogenetic diversity (16S OTUs) | 0.026 | NS |
| | $NO_3^-$-N ($mg \cdot kg^{-1}$) | Phylogenetic diversity (18S OTUs) | <0.001 | NS |

† NS represents not significant at 0.05.

**Table S5** ANOSIM analysis of the between-group differences of important indicator OTUs of three soil types.

| **16S OTUs** | | | | **18S OTUs** | | | |
|---|---|---|---|---|---|---|---|
| **Rodent** | **Group**[†] | ***R*** | ***p*** | **Rodent** | **Group** | ***R*** | ***p*** |
| Marmot | I vs O | 0.338 | 0.002 | Marmot | I vs O | 0.317 | 0.003 |
| | **A vs O** | **0.976** | **<0.001** | | **A vs O** | **0.864** | **<0.001** |
| | A vs I | 0.688 | <0.001 | | A vs I | 0.526 | <0.001 |
| Squirrel | I vs O | 0.738 | <0.001 | Squirrel | I vs O | 0.204 | 0.013 |
| | **A vs O** | **0.926** | **<0.001** | | **A vs O** | **0.708** | **<0.001** |
| | A vs I | 0.136 | NS[‡] | | A vs I | 0.432 | <0.001 |
| Gerbil | I vs O | 0.881 | <0.001 | Gerbil | I vs O | 0.822 | <0.001 |
| | **A vs O** | **0.924** | **<0.001** | | **A vs O** | **0.879** | **<0.001** |
| | A vs I | 0.103 | NS | | A vs I | 0.394 | <0.001 |
| Vole | I vs O | 0.733 | <0.001 | Vole | I vs O | 0.344 | 0.003 |
| | **A vs O** | **0.973** | **<0.001** | | **A vs O** | **0.869** | **<0.001** |
| | A vs I | 0.095 | NS | | A vs I | 0.724 | <0.001 |

[†] The group in bold indicates that this group has the largest ANOSIM statistic *R* value in comparison with other groups in terms of the given data and rodent species. A, I, and O represent active burrow, inactive burrow, and off-burrow soils, respectively.

[‡] NS represents not significant at 0.05.

**Table S6** ANOSIM analysis of the between-group abundance differences of important indicator bacterial OTUs co-occurring with amoebae at phylum level.

| **Data**† | **Group**‡ | ***R*** | ***p*** |
|---|---|---|---|
| Ind_A | I.Bur vs O.Bur | 0.149 | <0.001 |
| | A.Bur vs O.Bur | 0.227 | <0.001 |
| | **A.Bur vs I.Bur** | **0.021** | **NS**§ |
| Ind_I | I.Bur vs O.Bur | 0.156 | <0.001 |
| | A.Bur vs O.Bur | 0.359 | <0.001 |
| | **A.Bur vs I.Bur** | **0.074** | **NS** |
| Ind_C | I.Bur vs O.Bur | 0.134 | <0.001 |
| | A.Bur vs O.Bur | 0.181 | <0.001 |
| | **A.Bur vs I.Bur** | **-0.004** | **NS** |

† Ind_A, Ind_I, and Ind_C represent the amoeba-co-occurring bacteria identified as active burrow, inactive burrow, and off-burrow indicator OTUs, respectively.

‡ The group in bold indicates that this group has the smallest ANOSIM statistic *R* value in comparison with other groups in terms of the given data. A.Bur, I.Bur, and O.Bur represent active burrow, inactive burrow, and off-burrow soils, respectively.

§ NS represents not significant at 0.001.

**Table S7** Definitions of the abbreviations in Figure 3.

| **Other** | **Taxonomy** |
|---|---|
| Others1 | *Acidobacteriales, Gaiellales, Puia* |
| Others2 | *Antricoccus, Muribaculaceae, Oscillospiraceae, Pedobacter, Pseudarthrobacter, Pseudomonas litoralis* |
| Others3 | NS11−12, KF−JG30−B3 |
| Others4 | *Cytophagales, NS11−12 marine group, RB41* |
| Others5 | *Cytophagales, Microscillaceae, NS11−12 marine group, RB41, KF−JG30−B3, TRA3−20* |

**Table S8** References for determining microbial functional traits in Figure 3.

| Taxa | Associated functional traits | References |
|---|---|---|
| *Rhodanobacter* | Nitrogen cycle | (1) |
| *Ligilactobacillus* | Nitrogen cycle | (2) |
| *Beijerinckiaceae* | Nitrogen cycle | (3) |
| *Nitrospira* | Nitrogen cycle | (4) |
| *Brachybacterium* | Disease | (5-7) |
| *Staphylococcus vitulinus* | Disease | (8) |
| *Carnobacterium inhibens* | Disease | (9) |
| *Bacteroides* | Disease | (10, 11) |
| *Bergeyella* | Disease | (12) |
| *Ruminococcus* | Disease | (13-15) |
| *Micriamoeba* sp. | Disease | (16) |
| *Balamuthia mandrillaris* | Disease | (17) |
| *Vermamoeba vermiformis* | Disease | (18-20) |
| *Acanthamoeba* genotype T3 | Disease | (19) |
